# Expanding the scope of *Mycobacterium abscessus* reference strains to improve pulmonary disease modeling

**DOI:** 10.1101/2025.11.19.689313

**Authors:** Ruth A. Howe, Chavis Tabor, Chandra M. Panthi, Guoqing Cheng, Binayak Rimal, Evan Chen, Ngan Nguyen, Gyanu Lamichhane

## Abstract

*Mycobacterium abscessus* (*Mab*) pulmonary disease is an emerging clinical challenge, particularly among individuals with immunosuppression or underlying structural lung conditions. There are currently no FDA-approved therapies for *Mab* disease. Existing treatment strategies using repurposed drugs are prolonged, complex, and yield low cure rates (30–50%), underscoring the urgent need for more effective therapeutics. Developing new treatments requires preclinical disease models that faithfully replicate human disease, and the choice of *Mab* strain is a key determinant of model relevance. The commonly used reference strain, ATCC 19977, was isolated from a non-pulmonary source but became the default due to its early availability. To evaluate its relevance for pulmonary disease modeling, we compared ATCC 19977 with 15 clinical *Mab* isolates derived from lung infections across diverse regions of the United States. In both *in vitro* assays and a validated mouse lung infection model, ATCC 19977 behavior differed from the clinical isolates for key traits including rapid systemic dissemination, failure to develop robust lung granulomas, and early mortality. In contrast, clinical isolates demonstrated greater pulmonary tropism and reduced dissemination, with several producing robust lung pathology. Based on these findings, we propose a set of pulmonary clinical isolates representing the major *Mab* subspecies for use in lung infection research. These isolates more accurately recapitulate the pathological features of human *Mab* lung disease and are expected to enhance the translational value of future mechanistic and therapeutic studies.

**SUMMARY STATEMENT:** Using a mouse model, this study identified *Mycobacterium abscessus* clinical isolates whose infection profiles more closely resemble human pulmonary disease, establishing them as superior reference strains to ATCC 19977 for future translational and therapeutic research.

## INTRODUCTION

### Why Mycobacterium abscessus matters

Nontuberculous mycobacteria (NTM) are ubiquitous environmental organisms that cause opportunistic infections in both healthy and immunocompromised individuals (Falkinham, 2022; Johansen et al., 2020). Pulmonary NTM disease is a rising clinical concern, particularly for patients with immune suppression, structural lung damage such as bronchiectasis or chronic pulmonary obstructive disease, cystic fibrosis, and impaired mucociliary clearance. Its rising incidence exceeds what can be explained by improved detection alone, suggesting changes in exposure or a growing population of susceptible hosts (Cristancho-Rojas et al., 2024; Victoria et al., 2021).

Among NTMs, *Mycobacterium abscessus* (*Mab*) is the second most common cause of pulmonary disease after *M. avium* and is notoriously difficult to treat, earning it the nickname “the clinical nightmare” (Lopeman et al., 2019; Nessar et al., 2012). A recent study reclassified *Mycobacterium abscessus* as *Mycobacteroides abscessus* due to its genome being distinct from the core genomics profile of species belonging to the *Mycobacterium* genus (Gupta et al., 2018). *Mab* often persists in patients who have received multiple antibiotic treatments for other conditions, transitioning from a colonizing bystander to an active pathogen (Benwill and Wallace, 2014; Kwak et al., 2019). As a result, *Mab* infections are frequently antibiotic-experienced and require prolonged, multidrug regimens (Daley et al., 2020; Recchia et al., 2023). Yet, fewer than half of patients achieve culture conversion, and therapies are expensive, toxic, and poorly tolerated (Kwak et al., 2019; Park et al., 2017). The lack of any FDA-approved *Mab* treatment, coupled with poor clinical outcomes, underscores the urgent need for new, more effective and tolerable therapies.

### Why accurate disease models matter

Progress in understanding *Mab* biology and disease pathogenesis as well as therapeutic development all depend on preclinical models that accurately reproduce human disease. When laboratory models fail to mirror the clinical scenario, they can yield partial or misleading insights that impede our ability to accurately understand disease pathogenesis and develop treatments. A well-known example comes from *Escherichia coli*: strain K-12, isolated in 1922 and used extensively in molecular biology, lacks the virulence plasmids, pathogenicity islands, and secretion systems of disease-causing strains like enterohemorrhagic *E. coli* O157:H7 (Perna et al., 2001). Consequently, results from K-12 often fail to predict how pathogenic *E. coli* behave in humans. Similar limitations in laboratory models have challenged research in bacteriology, parasitology, mycology, oncology, neuroscience, and others, illustrating a key principle: model systems must reflect the biology of human disease. As fields mature, it is common for multiple models to be developed to reflect specific disease scenarios and improve therapeutic development.

For *Mab*, most laboratory studies have relied on the ATCC 19977 strain, originally isolated from a knee abscess in 1950 (Moore and Frerichs, 1953). Because ATCC 19977 was the first, and for many years the only *Mab* strain available from the American Type Culture Collection, it became the de facto reference strain for experimental and comparative studies. While its early adoption standardized research within the field, heavy reliance on this strain now limits progress. Most current *Mab* infections are pulmonary, raising concern that a strain derived from a knee abscess may not accurately model respiratory disease pathogenesis, host response, or drug efficacy (Dartois et al., 2024).

### Why *Mab* disease modeling is complicated

*Mab* disease can result from multiple routes of infection that lead to distinct disease outcomes. Direct inoculation through trauma or contaminated surgical equipment causes localized abscesses, as in the 1950 index case from which ATCC 19977 was derived (Moore and Frerichs, 1953). In contrast, pulmonary infections result primarily from inhalation of aerosolized *Mab* originating from biofilm reservoirs in domestic plumbing and aqueous environments (Norton et al., 2020; Ruis et al., 2021; Thomson et al., 2025; Virdi et al., 2021).

Both routes can occasionally lead to systemic dissemination, producing diverse bacterial subpopulations with differing drug susceptibilities (Park et al., 2023). Whether dissemination depends more on bacterial factors or host conditions remains uncertain. Historical descriptions of the 1950 case provide limited insight into systemic spread, and some apparent dissemination may have resulted from clinical practices of the time rather than hematogenous infection. The pulmonary versus inoculation infection routes expose *Mab* to very different host environments and evolutionary pressures. Direct inoculation produces aggressive, abscess-forming infections even in immunocompetent individuals, while pulmonary infection often begins as a chronic colonization that slowly progresses to granulomatous disease in immunocompromised hosts (Lee et al., 2015). It remains unclear whether a single environmental or abscess-derived strain can naturally cause both forms of disease without additional adaptation. Consequently, strains isolated from non-pulmonary sources may not faithfully represent pulmonary pathogenesis.

Adding further complexity, *Mab* comprises at least two dominant subspecies, *abscessus* and *massiliense*, that differ in drug susceptibility due to Erm(41) activity (Tortoli et al., 2016). A third, less common subspecies, *bolettii*, has not yet reached clinical prominence and additional environmental variants may exist. Differences in subspecies, colony morphotypes (smooth versus rough), and infection source (pulmonary versus extrapulmonary) all influence disease behavior and should be reflected in model selection (López-Roa et al., 2022).

### Significance of this study

We asked whether ATCC 19977, recovered from a knee abscess over 75 years ago, still adequately represents the *Mab* strains causing today’s pulmonary infections. To address this, we compared ATCC 19977 with *Mab* isolates obtained from patients with lung disease originating from distinct geographical locations, encompassing both major subspecies. Using *in vitro* assays and a validated mouse model based on aerosol infection, we identified strains that reproduced key features of human *Mab* pulmonary disease. These isolates more accurately reflect current clinical infections and a subset are proposed as improved reference strains for future pathogenesis and therapeutic studies.

## RESULTS

### Subspecies and Morphotype

Clinically isolated *Mab* strains representing subspecies *abscessus* and *massiliense,* obtained from the lungs of patients from distinct regions of the United States were included in this study (Table 1). One *in vitro* phenotype relevant to *Mab* pathogenesis is its ability to form biofilms in growth containers (Greendyke and Byrd, 2008). We hypothesized that the porosity of the growth substrate influences *Mab*’s biofilm-forming capacity. To test this, we cultured *Mab* isolates in tubes made of materials with differing densities and porosities. All growth media contained 0.05% Tween-80 to promote planktonic growth, and cultures were incubated under constant orbital shaking at 220 RPM. Under these conditions, true biofilm formers would be expected to overcome inhibition from Tween-80 to aggregate or attach to substratum.

**Table 1:** *Mycobacterium abscessus* pulmonary clinical isolates, subspecies information, sourcing, and growth phenotypes. Appearance on agar refers to colony appearance on agar (Figure S2). Appearance in broth refers to appearance of culture in test tubes (Figure S1). Composite morphotype refers to overall behavior on solid and in liquid media. Abbreviations: Johns Hopkins Hospital, Baltimore, Maryland (JHH), National Jewish Health, Denver, Colorado (NJH).

| Isolate ID | Subspecies | Source | Appearance on Agar | Appearance in Broth | Composite Morphotype |
| --- | --- | --- | --- | --- | --- |
| M9502 | *abscessus* | JHH | Mixed | Planktonic | Mixed |
| M9503 | *abscessus* | JHH | Smooth | Planktonic | Smooth |
| M9507 | *abscessus* | JHH | Smooth | Planktonic | Smooth |
| M9508 | *abscessus* | JHH | Rough | Planktonic, heavy aggregate, and heavy biofilm | Rough |
| M9510 | *massiliense* | JHH | Smooth | Planktonic | Smooth |
| M9533 | *abscessus* | JHH | Smooth | Planktonic | Smooth |
| M9624 | *abscessus* | NJH | Smooth | Planktonic | Smooth |
| M9626 | *abscessus* | NJH | Mixed | Planktonic, minimal aggregate, and biofilm | Mixed |
| M9643 | *abscessus* | NJH | Mixed | Planktonic | Mixed |
| M9647 | *massiliense* | NJH | Rough | Planktonic, heavy aggregate, and heavy biofilm | Rough |
| M9654 | *abscessus* | NJH | Smooth | Planktonic | Smooth |
| M9656 | *abscessus* | NJH | Smooth | Planktonic | Smooth |
| M9660 | *abscessus* | NJH | Mixed | Planktonic, heavy aggregate, and biofilm | Rough |
| M9672 | *abscessus* | NJH | Mixed | Planktonic and biofilm | Mixed |
| M9674 | *abscessus* | NJH | Smooth | Planktonic | Smooth |

In low-density, high-porosity materials such as polypropylene and polystyrene, *Mab* strains exhibited a range of growth behaviors, from forming prominent meniscal biofilms to entirely planktonic cultures (**Figure S1, Table 1**). Some strains also demonstrated a wall-climbing phenotype, establishing discrete colonies along the tube walls. In contrast, in borosilicate glass tubes, characterized by high density and low porosity, growth ranged from pelleted aggregates settling at the bottom of the tubes to fully planktonic forms dispersed homogeneously throughout culture broth. While variable meniscal films were observed, no stable biofilm adhered to the air–liquid interface or displayed climbing behavior.

Another *in vitro* phenotype relevant to disease pathogenesis is colony morphotype on agar media. *Mab* colonies are classically described as either “smooth,” featuring glabrous domes with circular margins, or “rough,” characterized by matte, irregular, and flaky colonies (Johansen et al., 2020). Although intermediate forms are not traditionally recognized, morphotype switching during infection has been documented (Byrd and Lyons, 1999; Howard et al., 2006). For ATCC 19977, colony morphology was reported as mostly smooth with rare rough variants (Moore and Frerichs, 1953) and maintained that variable morphology in our hands. In contrast, clinical isolates consistently displayed uniform colony morphologies across a plate but rarely matched the classical extremes. Most exhibited smooth or mixed characteristics (**Figure S2**). For clarity and comparison within the traditional smooth–rough framework, we categorized each strain as smooth, rough, or mixed based on its collective behavior in both liquid and solid media (**Table 1**).

### Drug Susceptibilities

Antibiotics can serve not only as therapeutic agents but also as insightful probes into *Mab* physiology. Because the action of an antibiotic depends on numerous cellular processes, such as metabolism, stress response, and cell envelope integrity, its minimum inhibitory concentration (MIC) reflects the *Mab*’s overall physiological state and its ability to cope with specific types of stress. In this way, MIC values provide an integrated snapshot of how different isolates respond to diverse perturbations. By systematically measuring MICs across multiple antibiotics, we can use these antibiotics as probes to classify isolates according to their physiological behavior. Grouping isolates based on shared MIC profiles allows us to identify those that represent a “consensus response,” serving as representative strains for *Mab*.

To this end, we determined the MICs of 24 antibiotics spanning agents currently used in clinical treatment and compounds in preclinical development against all *Mab* isolates included in this study (**Table S1**). As anticipated for clinical strains collected at different stages of infection and treatment, MICs for commonly used antibiotics varied considerably. Many clinical isolates exhibited resistance to macrolides azithromycin and clarithromycin, and a subset displayed near pan-resistance to multiple agents. These MIC patterns not only highlight the phenotypic diversity of *Mab* but also offer a functional framework for probing the underlying physiological and adaptive mechanisms that shape antibiotic susceptibility. *Mab* is notorious for discordance between *in vitro* MIC and *in vivo* efficacy, which is postulated to be mediated by both limitations from slow growth relative to antimicrobial degradation *in vitro* and *in vivo* organization into protective biofilms or cavitations (van Ingen and Kuijper, 2014). This poor concordance underlies the Clinical and Laboratory Standards Institute (CLSI) recommendation against performing MICs for non-macrolides to inform *Mab* treatment. In this study, strain morphotypes of smooth, rough, or mixed did not directly correlate with antibiotic susceptibilities *in vitro* for any agent. However, the correspondence of these morphotypes to biofilm formation or other structural organization in *vivo* may impact strain susceptibility to these agents and will require additional study.

### *In vivo* characteristics of ATCC 19977

A mouse model of *Mab* infection provides a controlled and tractable system for assessing the virulence potential of clinical isolates (Dartois et al., 2024). In this context, virulence is defined by the ability of an isolate to establish and sustain infection in the lungs, trigger host immune responses, disseminate to other organs, and induce pathological changes. Because these outcomes reflect the combined effects of numerous *Mab* determinants interacting with host defenses, the mouse model captures an integrated biological measure of pathogenic potential. By comparing these parameters using an identical protocol with *Mab* isolates as the only variable, distinct virulence patterns can be discerned, enabling the identification of a representative *Mab* strain whose infection dynamics, proliferation, and pathology profiles typify those produced by most clinical isolates and mirror the disease characteristics observed in human *Mab* infections.

We used a validated mouse model of *Mab* lung infection (Galanis et al., 2022; Maggioncalda et al., 2020) to characterize *in vivo* growth and dissemination of the isolates. To assess the chronic phase of infection, we extended the infection period to eight weeks. C3HeBFe/J (Kramnik) mice were infected with aerosolized ATCC 19977 and subjected to dexamethasone-based immunosuppression regimens as described in the original model protocol (Maggioncalda et al., 2020) (**Figure 1A**). While a daily dose of 8 mg/kg dexamethasone was sufficient to sustain infection during the early phase, this dose failed to maintain or promote *Mab* persistence and growth, particularly for clinical isolates, through the eight-week period. Therefore, the dexamethasone dose was increased to 10 mg/kg after week two post-infection to support extended infection. Under a regimen of 10 mg/kg/day dexamethasone, ATCC 19977 clearance from the lungs began at four weeks post-infection, with complete clearance observed after immunosuppression withdrawal at six weeks (8+10+0 regimen; **Figure 1B–D**) prompting adjustment of dexamethasone dosing. Increasing dexamethasone dose to 12 mg/kg/day at six weeks (8+10+12 regimen) allowed ATCC 19977 to maintain a higher lung bacterial burden with stable dissemination to the spleen and liver.

**Figure 1.**
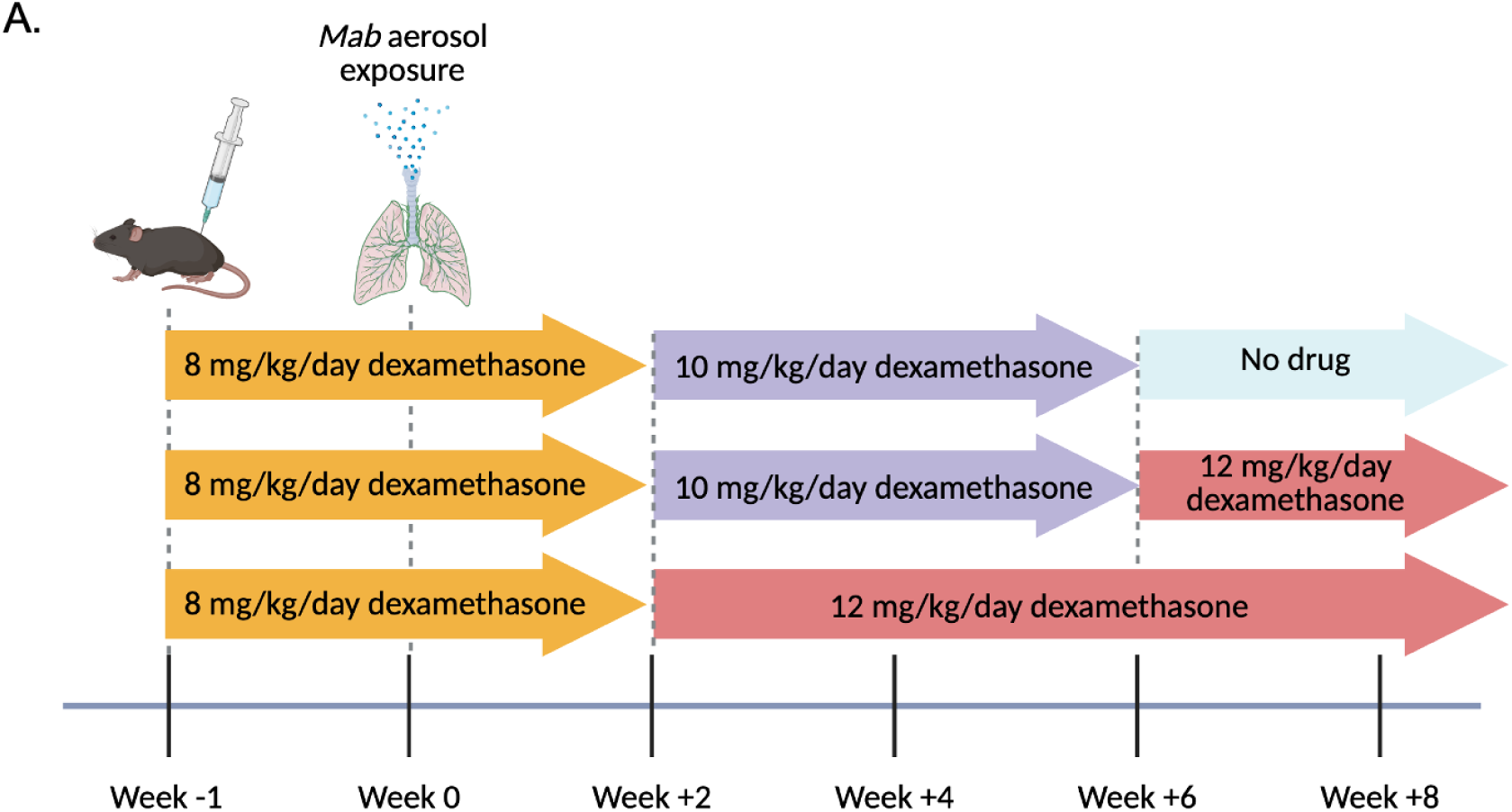
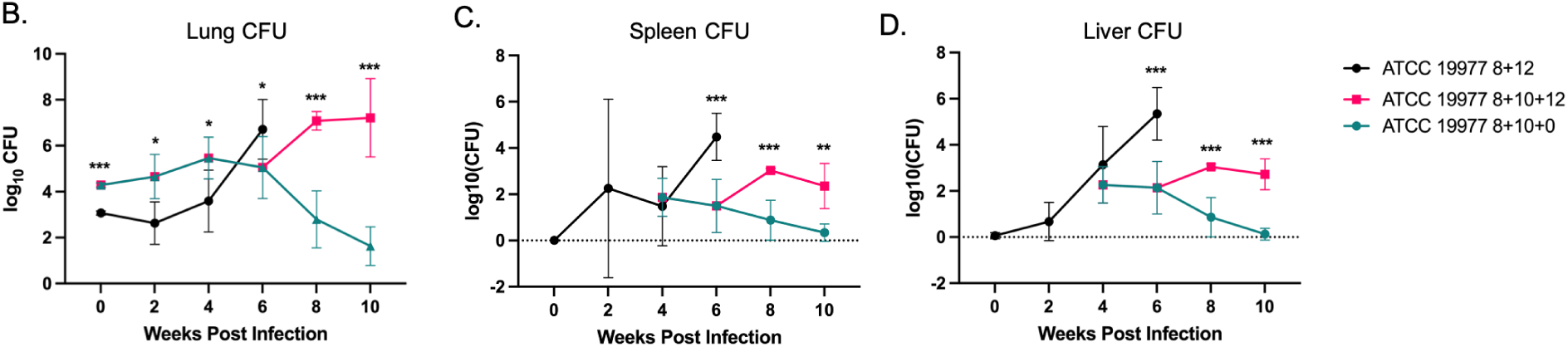
A. Schematic of immunosuppression and infection used to determine activity of ATCC 19977 reference strain in Kramnik mice. Figure created with BioRender. B-D *Mycobacterium abscessus* burden as determined by colony forming unit (CFU) assay in B. lung, C. liver, and D. spleen in mice infected with ATCC 19977 under different immunosuppressive schemes with dexamethasone (dex). Teal triangles indicate mice receiving 3 weeks of 8 mg/kg/day dex followed by 4 weeks of 10 mg/kg/day dex. Pink squares indicate mice receiving 3 weeks of 8mg/kg/day dex followed by 4 weeks of 10 mg/kg/day dex, followed by 12 mg/kg/day dex until death. Black circles indicate animals receiving 3 weeks of 8 mg/kg/day dex followed by 12 mg/kg/day dex until death.

However, clinical *Mab* isolates could not sustain pulmonary infection under either the 8+10+0 or 8+10+12 dexamethasone regimen (data not shown). These isolates required a more intensive immunosuppression schedule—8 mg/kg/day dexamethasone for the first two weeks post-infection followed immediately by 12 mg/kg/day (8+12 regimen)—to maintain or increase pulmonary bacterial loads (**Figure 2**). To provide a valid comparator, we also profiled ATCC 19977 using the same 8+12 regimen (**Fig. 1B–D**). Under these conditions, ATCC 19977 caused early and aggressive systemic dissemination. Animals with lower pulmonary bacterial burdens succumbed early, exhibiting widespread microabscesses upon necropsy. All animals died before the eight-week endpoint, and histopathological analysis revealed *Mab*-positive lesions or scarring in only 50% of lungs examined (**Table 2**), suggesting that mortality was frequently due to extrapulmonary complications rather than primary pulmonary pathology.

**Figure 2.**
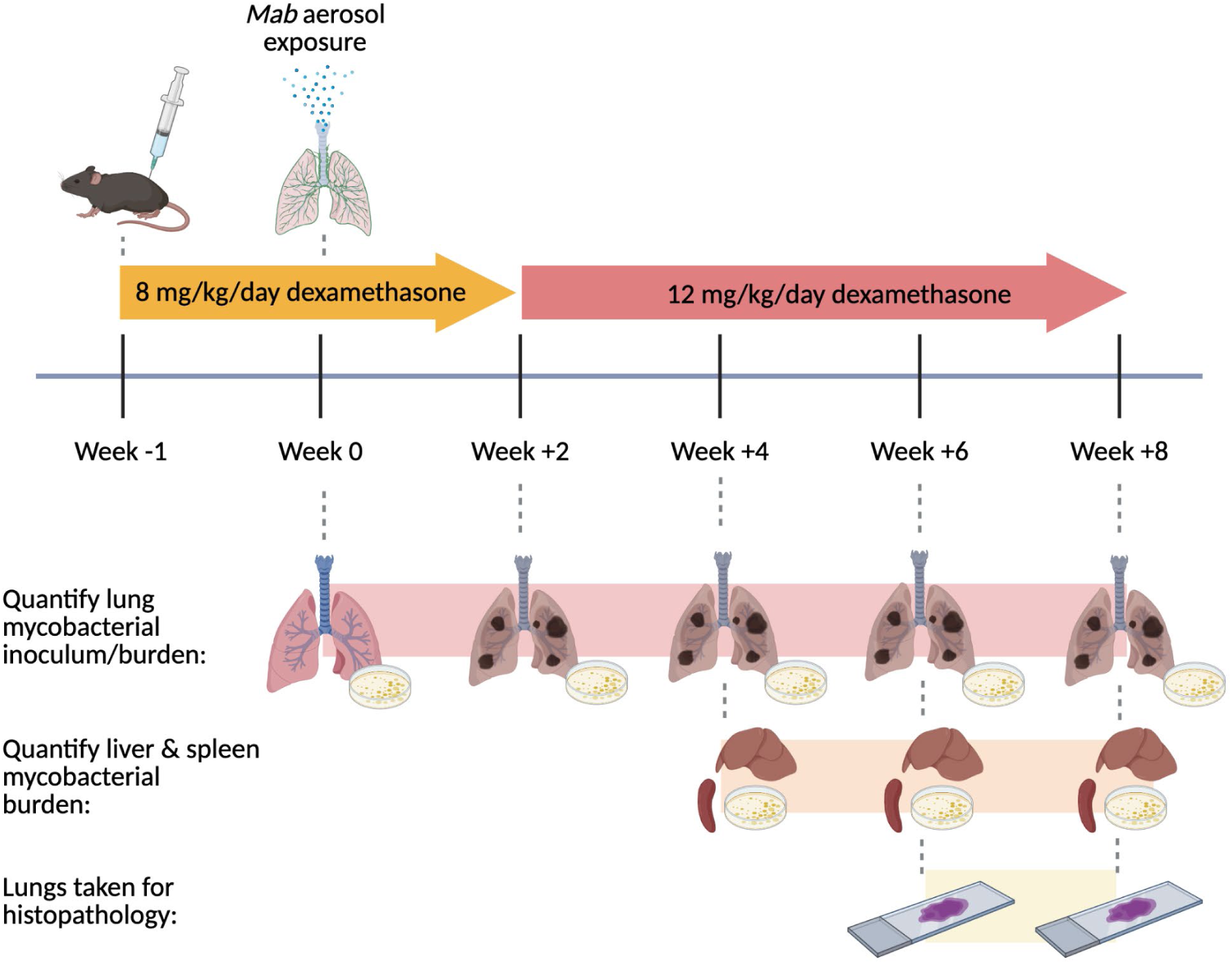
Schematic for immunosuppression, infection, and analysis for C3HeB/FeJ mice infected with ATCC 19977 and clinical pulmonary *Mab* isolates.

**Table 2.** Lung Histopathology findings at 0-6 weeks post infection for ATCC 19977-infected mice receiving 8mg/kg/day dexamethasone until 2 weeks post infection and 12 mg/kg/day dexamethasone thereafter.

| Weeks post infection | Lung | Spleen | Liver |
| --- | --- | --- | --- |
| 0 | Normal | Normal | Normal |
| 2 | Normal structure, but small AFB focus without robust inflammatory response in one specimen | Normal | Normal |
| 4 | Tissue invasion of AFB seen in 50% samples without robust immune response. Overall structure preserved in all specimens. | Normal | 50% samples with severe steatosis, no AFB organisms seen |
| 6 | Diffuse invasion of lung spaces by AFB organisms with purulence in bronchioles in 50% of samples, no organizing granulomas seen. Gross structure preserved in all specimens | 50% samples with splenomegaly and areas of scarring. No discrete AFB seen in any specimen. | Some biliary sludge & congestion, but no AFB+ organisms seen. |
| 7 (death) | Large AFB+ granuloma in one specimen with diffuse AFB tissue invasion in other specimen | Splenomegaly with occasional fuchsin-positive subcapsular cells, but no discrete AFB seen. | Steatosis with one sample showing AFB+ microabscesses |

### *In vivo* characteristics of pulmonary clinical *Mab* isolates

#### Lung-derived isolates produced greater pulmonary disease burden than ATCC 19977

All lung-derived clinical *Mab* isolates established persistent infections in the mouse lung, although the magnitude and rate of proliferation differed among isolates as determined by CFU enumeration (**Figure 3**). For most isolates belonging to both subspecies *abscessus* and *massiliense*, pulmonary burdens remained elevated during the early stages of infection and persisted through the 8-week endpoint, with all mice surviving to this final time point. Clinical isolates M9647 and M9660 were the exceptions, with all animals dying before the 8-week timepoint. The severity of structural lung damage also varied across isolates and generally correlated with the extent of *Mab* expansion (**Table 3**).

**Figure 3.**
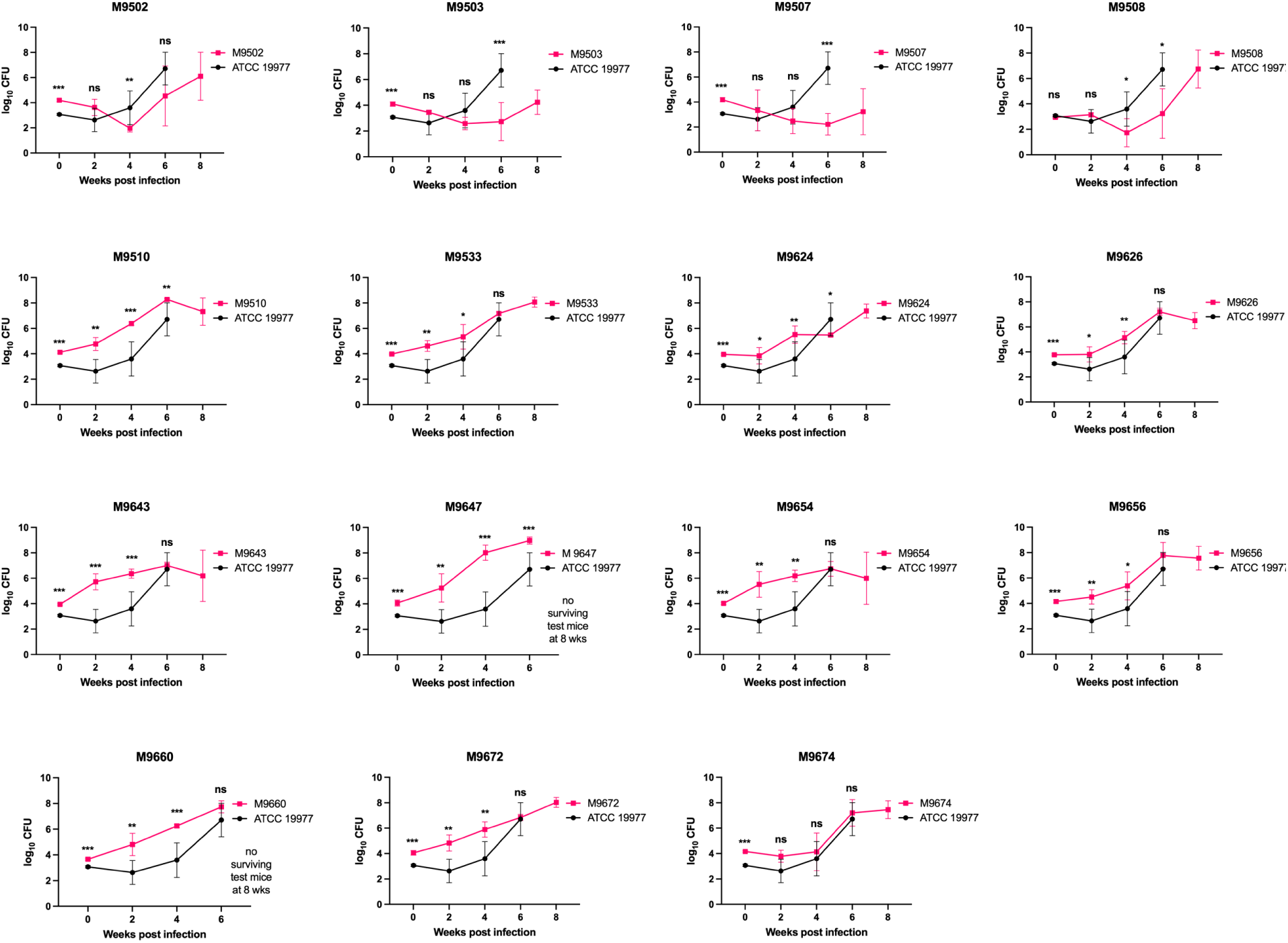
Lung CFU burden of pulmonary-derived *Mab* isolates (pink) compared to ATCC 19977 (black). Statistical significance is reported as p-value with *= <0.05, **=<0.01, ***=<0.001 or ns denoting nonsignificant.

**Table 3.** Lung histopathology findings at weeks 6-8 post infection for mice infected with clinical *Mab* isolates. The two subspecies *massiliense* isolates are indicated with * and the remaining isolates are subspecies *abscessus*.

| Clinical Isolate ID | Timepoints analyzed | Lung Histopathology findings | Comparison to ATCC19977 lung pathology |
| --- | --- | --- | --- |
| M9502 | Week 8, 10 | Neither lung at week 8 with visible structural lung disease. At week 10, one of two samples with large AFB+ granuloma | **Less structural lung disease** at week 8 than for ATCC 19977, but comparable space-occupying lesions and structural damage at week 10 of M9502 as at week 6-7 of ATCC 19977. |
| M9503 | Week 8, 10 | No AFB organisms seen at any timepoint and no structural lung disease identified | **Less structural lung disease** compared to ATCC 19977. |
| M9507 | Week 6, 8, 10 | No AFB organisms or structural lung disease seen at any timepoint | **Less structural lung disease** compared to ATCC 19977. |
| M9508 | Week 6, 8, 10 | Extensive AFB+ granulomatous disease at all timepoints | **Substantially increased structural lung disease** at weeks 8 and 10, with comparable disease at week 6 to ATCC 19977 |
| M9510* | Week 6, 8 | At week 6, large-volume AFB+ granulomas seen in 2/2 samples. Mice that survive to week 8 have some scarring, but no remaining AFB+ organisms seen | **Substantially increased structural lung disease** at week 6 than with ATCC 19977. Mice surviving to 8 weeks appear to have controlled disease but display evidence of prior infection. |
| M9533 | Week 6, 8 | In ½ samples for both timepoints, see variably-sized granulomatous disease | **Comparable lung disease** burden at week 6 of infection to ATCC 19977, with longer survival but less penetrance of lung disease at week 8. |
| M9624 | Week 6, 8 | One of two samples at 6 weeks with single AFB+ lesion surrounding bronchus with invasion through mucosa. Other lung with no clear structural disease or AFB at 6 weeks. No AFB or scarring at 8 weeks in either sample. | **Less structural damage** and lower visible AFB burden at 6 week timepoint than for ATCC 19977. Mice surviving to 8 weeks appear to have controlled pulmonary disease. |
| M9626 | Week 6, 8 | Multiple large AFB+ granulomas in addition to bronchial AFB+ debris, perivascular invasion, and free AFB. | **Substantially more structural damage** to lungs at 6-week timepoint than for ATCC 19977 and with continued progression at 8 week timepoint. |
| M9643 | Week 6, 8 | For each timepoint, extensive AFB+ granulomas in ½ samples, but no AFB organisms or scarring/structural disruption seen in other sample. | **Comparable structural damage** to lungs in some mice at 6-week timepoint, but with same pattern persisting at 8 weeks, showing less penetrance at that timepoint than for ATCC 19977. |
| M9647* | Week 6 | Widespread, high-burden AFB+ granulomas (variable sizes) with areas of hemorrhage, invasion into airways, and substantial sclerosis in both samples for week 6 samples. No animals surviving at week 8 for analysis. | **Substantially more structural damage** to lungs at the 6-week timepoint with M9647 |
| M9654 | Week 6, 8 | Moderate to High-AFB burden granulomas throughout lungs, with more structural damage seen at the 6 week timepoint than at the 8 week timepoint. | **Substantially more structural damage** to lungs at the 6-week timepoint with M9654. *\*\*mice that survive to 8 weeks begin to clear disease* |
| M9656 | Week 6, 8 | At week 6, ½ lungs with solitary small focus of AFB+ granulomatous change. Lungs otherwise normal. No AFB seen at 8 weeks and only one small area of sclerosis. | **Less structural damage** and lower visible AFB burden at 6 weeks compared to ATCC 19977 and disease resolving in animals surviving to week 8. |
| M9660 | Week 6 | Peribronchial & mucosal AFB invasion in one specimen with paucibacillary granulomas in second specimen. | **Comparable to slightly increased structural lung disease** at week 6 compared to ATCC 19977. |
| M9672 | Week 6, 8 | At week 6, ½ lungs with single moderate-sized AFB+ granuloma in otherwise structurally normal lung. No evidence of structural disease at 8 weeks | **Comparable to less structural lung disease** at week 6 compared with ATCC 19977 and animals surviving to week 8 showed resolving disease.. |
| M9674 | Weeks 6, 8 | ½ specimens at 6 weeks and 2/2 specimens at 8 weeks with high-AFB-burden peribronchial and airway infiltration without organization into granulomas. Do see mats of AFB in airways | **Comparable AFB disease burden** with possibly less granuloma formation and <u>more biofilm formation</u> in M9674 at 6 and 8 weeks. |

Isolates M9502, M9503, and M9507 produced low but stable lung burdens and showed no histopathologic evidence of lung injury. These characteristics suggest that such isolates can maintain chronic infections over extended periods and may represent useful models for studying chronic *Mab* pulmonary infection with minimal tissue pathology. In contrast, several isolates produced more pronounced disease. Compared with the reference strain ATCC 19977, certain isolates displayed comparable levels of lung pathology by 6–8 weeks post-infection. However, subspecies *abscessus* isolates M9508, M9626, M9654, and M9674, along with subspecies *massiliense* isolates M9510 and M9647, induced more extensive structural lung damage and sustained higher burdens at terminal time points. The lung burdens and pathology produced by these isolates more closely reflected those observed in patients with chronic *Mab* pulmonary disease (**Table 3**).

#### Lung-derived isolates exhibit reduced systemic dissemination compared to ATCC 19977

Infection with ATCC 19977 resulted in the rapid onset of pulmonary lesions containing abundant acid-fast bacilli (AFB) as early as two weeks post-infection. By four weeks post infection, these lesions showed evidence of tissue invasion within the lungs, accompanied by visible hepatic abnormalities, and by six weeks, the mice exhibited pronounced splenomegaly. CFU quantification confirmed systemic dissemination, with *Mab* burdens reaching approximately 10⁴ CFU in the spleen and 10⁵ CFU in the liver at six weeks (**Figures 1B, 4, and 5**). In contrast, mice infected with lung-derived clinical isolates displayed markedly reduced and delayed dissemination to both spleen and liver (**Figures 4 and 5**). Even among isolates associated with early mortality, such as M9647 and M9660, systemic bacterial loads were substantially lower—exceeding a 1 log₁₀ reduction in the spleen and more than a 2 log₁₀ reduction in the liver compared with ATCC 19977 at the same 6-week time point. These findings suggest that, unlike ATCC 19977, which readily disseminates beyond the lungs, pulmonary-derived clinical isolates primarily cause localized disease confined to the lungs. Consequently, mortality in these infections is likely driven by progressive pulmonary pathology rather than by systemic *Mab* spread.

**Figure 4.**
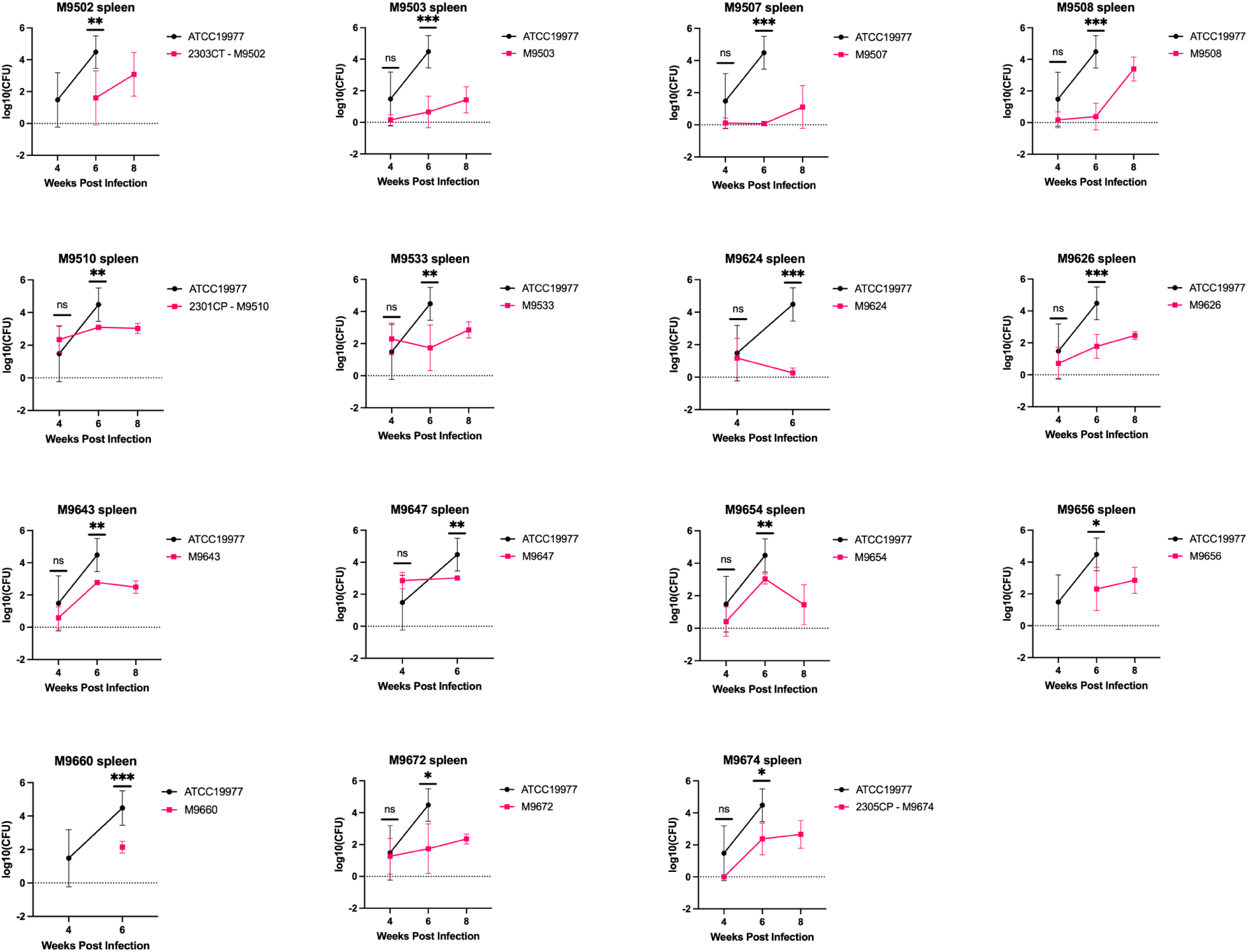
Spleen CFU burden of pulmonary-derived *Mab* isolates (pink) compared to ATCC 19977 (black). Statistical significance is reported as p-value with *= <0.05, **=<0.01, ***=<0.001 or ns denoting nonsignificant.

**Figure 5.**
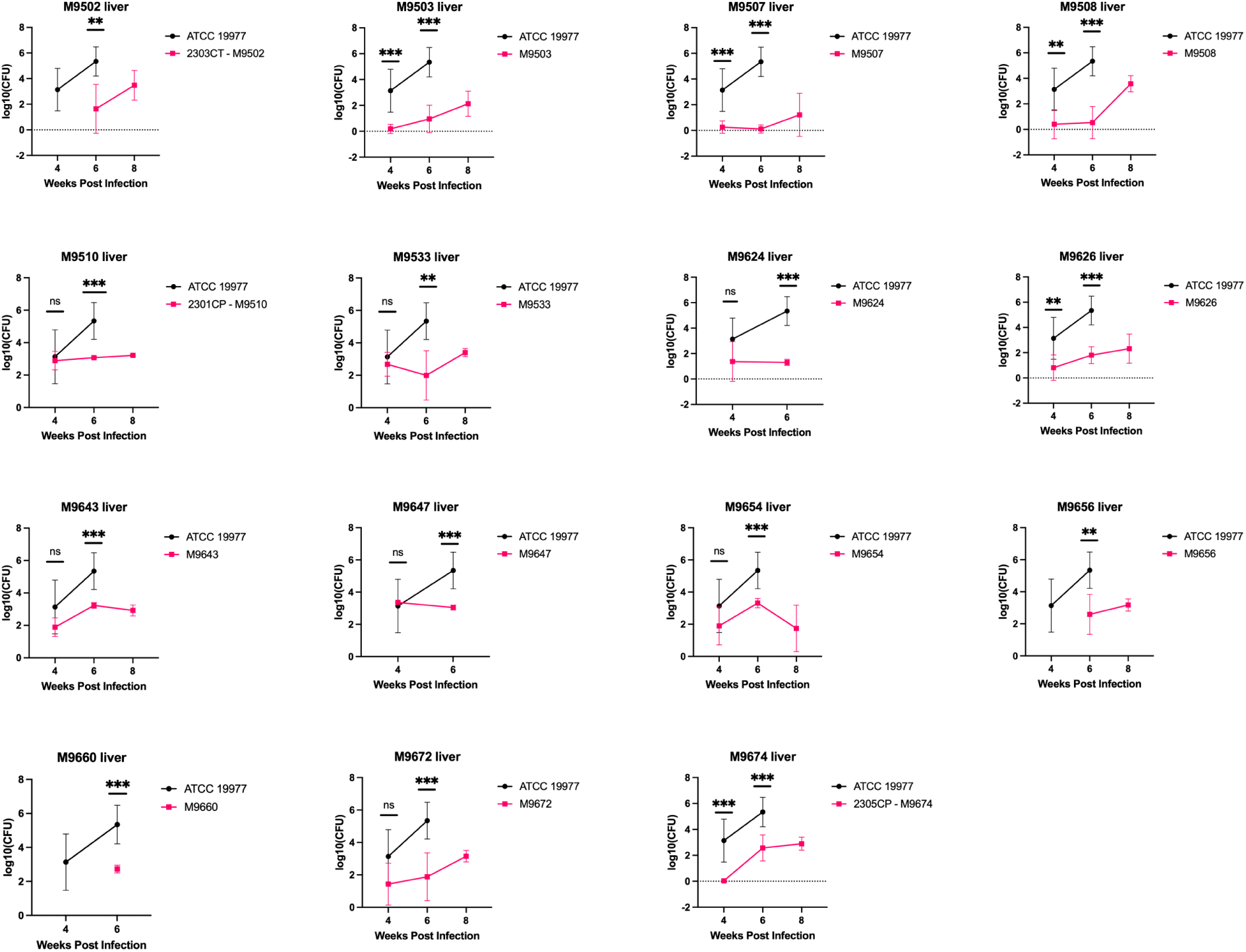
Liver CFU burden of pulmonary-derived *M. abscessus* isolates (pink) compared to ATCC 19977 (black). Statistical significance is reported as p-value with *= <0.05, **=<0.01, ***=<0.001 or ns denoting nonsignificant.

### Proposed *Mab* reference strains

Based on comparative analyses of their *in vivo* lung growth profiles, associated histopathologic changes, dissemination patterns to extrapulmonary organs, colony morphotypes, and antibiotic susceptibility (MIC) profiles, five *Mab* isolates emerged as compelling candidates to serve as new reference strains (**Table 4**). Collectively, these isolates represent both major *Mab* subspecies (*abscessus* and *massiliense*) and encompass the diversity of colony morphotypes (smooth, rough, and mixed) and culture phenotype (planktonic and biofilm) as well as the spectrum of virulence observed among clinical isolates. They were therefore selected as representative of contemporary *Mab* strains whose biological and pathogenic characteristics more closely reflect those encountered in current patient populations.

**Table 4.** Summary of recommended strains for high-penetrance modeling of *Mab abscessus* and *Mab massiliense* pulmonary disease and primary characteristics. Composite Morphotype refers to combined behavior in liquid and on solid media (Table 1). Abbreviations: CFU colony forming unit, CLR clarithromycin, MIC minimum inhibitory concentration.

| Isolate ID | Subspecies | Composite Morphotype | Lung CFU burden compared to ATCC 19977 | Disease penetrance by 8 weeks | Systemic dissemination | Median CLR MIC (µg/mL) |
| --- | --- | --- | --- | --- | --- | --- |
| M9508 | *abscessus* | Rough | Higher burden, delayed onset | 100% | Some, late in course | 24 |
| M9626 | *abscessus* | Mixed | Higher burden | 100% | Some, late in course | 24 |
| M9654 | *abscessus* | Smooth | Higher burden | 100% | Some, late in course | 20 |
| M9510 | *massiliense* | Smooth | Higher burden | 100% | Some, late in course | 4 |
| M9647 | *massiliense* | Rough | Higher burden | 100% | Some, late in course | 10 |

Adopting these isolates as reference strains in future *Mab* studies is expected to produce data that are more clinically relevant than those generated using the long-established laboratory strain ATCC 19977. Because these isolates better reproduce the infection dynamics, pathology, and drug susceptibility patterns seen in human *Mab* disease, experimental outcomes derived from them are likely to have greater translational value and applicability to real-world clinical infections.

We propose three reference strains for *Mab* subspecies *abscessus* and two for subspecies *massiliense*, each chosen for its consistent ability to induce reproducible lung pathology and to model distinct clinical disease states.

### *Mab* subspecies *abscessus* M9508 (rough morphotype on agar, and planktonic on broth)

This strain produces progressive lung disease characterized by structural tissue damage evident by eight weeks post-infection, with animals surviving through 10 weeks and showing late-stage systemic dissemination. These features make M9508 well suited for modeling chronic *Mab* pulmonary disease with rough-strain conversion and secondary dissemination, as seen in older cystic fibrosis patients with advanced infection.

### *Mab* subspecies *abscessus* M9626 (mixed morphotype on agar, and planktonic and biofilm growth on broth)

M9626 causes a more rapid onset of pulmonary disease compared to M9508 but exhibits limited extrapulmonary spread. Its faster disease progression and primarily lung-confined pathology make it a valuable model for studying acute or rapidly progressive infections, such as those occurring in transplant recipients, and for evaluating inhaled therapeutic interventions where systemic dissemination is minimal.

### *Mab* subspecies *abscessus* M9654 (smooth morphotype on agar and planktonic on broth)

This isolate shows a disease course similar to M9626 but has slightly lower macrolide MICs. Because of this characteristic, M9654 is particularly useful for testing macrolide-based combination therapies or regimens targeting early or less disseminated forms of *Mab* lung disease.

### *Mab* subspecies *massiliense* M9510 (smooth morphotype on agar and planktonic on broth)

Consistent with the massiliense subspecies, M9510 exhibits low macrolide MICs associated with an inactive *erm(41)* gene. It produces high pulmonary bacterial burdens but minimal dissemination, making it an excellent model for localized, non-disseminating lung infections dominated by smooth morphotypes.

### *Mab* subspecies *massiliense* M9647 (rough morphotype on agar, and planktonic and biofilm growth on broth)

Like M9510, M9647 maintains high pulmonary bacterial loads with limited systemic spread but exhibits the rough colony morphology associated with enhanced tissue invasion and inflammation. This isolate models rough morphotype-driven pulmonary disease that progresses more rapidly but remains primarily lung-localized. Though it also has a non-functional *erm(41)* gene, the macrolide MICs were slightly higher than for M9510, possibly because of its extensive biofilm formation.

Both *massiliense* strains (M9510 and M9647) demonstrate faster pulmonary disease progression with less systemic dissemination than the *abscessus* isolates, making them suitable for evaluating both inhaled and systemic (oral or injectable) therapeutic regimens. Together, these five isolates form a robust toolkit for modeling the breadth of *Mab* pulmonary disease observed in patients and for advancing the preclinical evaluation of novel anti-*Mab* therapies.

## DISCUSSION

Reference strains that accurately represent contemporary *Mab* isolates from pulmonary infections are essential for advancing both fundamental and translational research. Most current *Mab* studies rely on the long-established laboratory strain ATCC 19977, which was originally isolated from a knee abscess in 1950 (Moore and Frerichs, 1953). Although this strain has been instrumental in early *Mab* research, owing to its availability, complete genome sequence, and well-characterized *in vitro* and *in vivo* properties, it does not represent the biological or clinical features of modern pulmonary isolates that now dominate patient infections.

Our findings demonstrate that ATCC 19977 differs markedly from contemporary lung-derived clinical isolates in terms of virulence, disease localization, and antibiotic susceptibility profiles. Whereas pulmonary *Mab* disease typically arises via aerosol exposure, involves complex host-pathogen interactions within lung tissue, and manifests in diverse morphotypes and subspecies, ATCC 19977 exhibits disease behavior more consistent with acute, systemic infection. In mice, ATCC 19977 caused extensive early extrapulmonary dissemination, leading to high bacterial burdens outside the lungs and significant mortality attributable to non-granulomatous, systemic disease. Within the lung, ATCC 19977 did not produce uniform pathology even at the later timepoints. These findings indicate that ATCC 19977 provides a poor model for subacute or chronic pulmonary infection, the clinical scenario most relevant to human disease. Consequently, preclinical studies that continue to rely exclusively on this strain may produce results that do not accurately predict treatment responses or disease outcomes in patients.

Over the past several decades, clinical and genomic studies have revealed extensive heterogeneity among *Mab* isolates, including differences in colony morphotypes (smooth versus rough), virulence, and subspecies (notably *Mab* subsp. *abscessus* and *Mab* subsp. *massiliense*) (Bohr et al., 2021; Gutiérrez et al., 2021; Yang et al., 2025). Each of these traits contributes to the wide range of disease phenotypes seen in patients and influences treatment response and persistence within the lung. Given this diversity, a single reference strain such as ATCC 19977 is insufficient to capture the full biological and clinical spectrum of *Mab* pulmonary disease.

To address this gap, our study identified a panel of lung-derived, clinically relevant *Mab* isolates that more faithfully reproduce pulmonary disease characteristics observed in patients. We propose that *Mab* subspecies *abscessus* strains M9508, M9626, and M9654, along with *Mab* subspecies *massiliense* strains M9510 and M9647, together constitute a more representative and tractable reference set for modeling *Mab* pulmonary infection (Table 4). These strains collectively encompass both major subspecies and include smooth, rough, and mixed morphotypes. Importantly, each demonstrated robust lung colonization and high pulmonary disease penetrance in mice, confirmed by both histopathology and lung CFU enumeration, while exhibiting minimal and delayed systemic dissemination compared to ATCC 19977.

Using these strains in preclinical research offers several advantages. Their high and consistent lung disease penetrance enables efficient experimental design, reducing the number of animals and resources needed to achieve meaningful results. Moreover, the limited extrapulmonary spread minimizes confounding mortality from systemic infection, thereby improving the interpretability of survival and treatment efficacy data in therapeutic studies, particularly those evaluating inhaled or lung-targeted interventions.

Although our present work focused on establishing these strains within a standardized murine infection model to directly compare their pathogenic behaviors, this reference set can be adapted for use in diverse experimental contexts. For example, they could be integrated into immunocompetent or immunocompromised mouse models, aerosol infection systems, or studies exploring host immune responses and drug pharmacodynamics. Ultimately, adopting this broader, biologically relevant panel of *Mab* reference strains will enhance the translational relevance of preclinical studies and accelerate progress toward effective therapies for this increasingly prevalent and difficult-to-treat respiratory pathogen.

## MATERIALS AND METHODS

### Ethics

Mouse handling and procedures were undertaken in adherence to the national and the Animal Care and Use guidelines approved by the Johns Hopkins University Animal Care and Use Committee (protocol number MO23M123).

### Bacterial strains, culture media, drugs and growth conditions

*Mycobacterium abscessus* (*Mab*) strain ATCC 19977, an isolate obtained from knee abscessus (Moore and Frerichs, 1953), was procured in 2014 from American Type Culture Collection (Manassas, VA). It was authenticated by genome sequencing and confirmed as belonging to the subspecies *abscessus* (Story-Roller et al., 2021). *Mab* isolates M9502, M9503, M9507, M9508, M9510 and M9533 were isolated from lung expectorates obtained from bronchiectasis and cystic fibrosis patients at the Johns Hospital (Baltimore, MD) between 2004 and 2020 and archived by the Johns Hopkins Clinical Mycobacteriology Laboratory. Similarly, *Mab* isolates M9624, M9626, M9643, M9647, M9654, M9656, M9660, M9672 and M9674 were obtained from lung samples from cystic fibrosis and non-cystic fibrosis patients at National Jewish Health (Denver, CO) between 2019 and 2021 and archived by the Clinical Mycobacteriology Laboratory. For all isolates, their genomes were sequenced and this data was used to determine their subspecies classification as described (Story-Roller et al., 2021).

All *Mab* isolates were inoculated from early-passage frozen stocks and cultured in Middlebrook 7H9 broth (Difco, catalog no. 271310) supplemented with 0.5% glycerol, 10% albumin-dextrose-salt enrichment, and 0.05% Tween-80 at 37°C in an orbital shaker at 220 RPM as described (Story-Roller et al., 2021) . For determining minimum inhibitory concentrations of drugs, Middlebrook 7H9 broth supplemented with 0.5% glycerol and 10% albumin-dextrose-salt enrichment without Tween-80 was used in U-bottom 96-well plates with 350 μL capacity per well.

Powdered form of drugs was purchased or donated from commercial vendors as follows: amikacin (Sigma-Aldrich, catalog no. A3650), amoxicillin (Sigma-Aldrich, catalog no. A8523), azithromycin (Sigma-Aldrich, catalog no. 75199), bedaquiline (MedChemExpress, catalog no. HY-14881A), cefdinir (VWR, catalog no. TCC3111), cefditoren (Sigma-Aldrich, catalog no. SML3624), cefoxitin (Sigma-Aldrich, catalog no. C4786), ceftazidime (Sigma-Aldrich, catalog no. C3809), cefuroxime (Sigma-Aldrich, catalog no. C4417) clarithromycin (Sigma-Aldrich, catalog no. C9742), clofazimine (Sigma-Aldrich, catalog no. C8895), dicloxacillin (Sigma-Aldrich, catalog no. D9016), doripenem (Sigma-Aldrich, catalog no. SML1220), faropenem (Octagon Chemicals), imipenem (Sigma-Aldrich, catalog no. I0160), linezolid (Octagon Chemicals), metronidazole (Sigma-Aldrich, catalog no. M3761), moxifloxacin (Sigma-Aldrich, catalog no. PHR1542), MRX-6038 (MicuRx Pharmaceuticals), omadacycline (Paratek Pharmaceuticals), oxacillin (Sigma-Aldrich, catalog no. 28221), penicillin V (Sigma-Aldrich, catalog no. PHR2644), rifabutin (Sigma-Aldrich, catalog no. R3530) and tebipenem (Octagon Chemicals).

### Smooth and rough colony morphotyping

*Mab* isolates were cultured under standardized liquid and solid growth conditions to ensure consistent morphological assessment. For liquid culture, isolates were inoculated into Middlebrook 7H9 broth supplemented with 0.2% glycerol, 10% albumin-dextrose-catalase (ADC), and 0.05% Tween 80 to minimize clumping. Cultures were incubated at 37 °C under aerobic, shaking conditions (220 rpm) in test tubes measuring 1–1.5 cm in internal diameter. Tubes composed of different materials (borosilicate glass, polystyrene, polypropylene, or resin) were used to evaluate planktonic or clumping growth behavior and formation of meniscal biofilms with or without wall-climbing behavior.

For smooth versus rough colony phenotyping, *Mab* cultures from mid-log phase in planktonic form were streaked onto Middlebrook 7H10 agar supplemented with 0.5% glycerol and 10% ADC. Plates were incubated at 37 °C for 3–5 days, or until discrete, well-isolated colonies were visible. Colony morphology was then assessed both macroscopically and microscopically. Visual inspection was performed under front- and back-lighting conditions, and images recorded using an Interscience Scan 300 imager. Morphotypes on solid media were categorized as follows. Smooth morphotype: colonies appeared glabrous, dome-shaped, and circular, with a smooth, moist, or oily surface sheen. Rough morphotype: colonies exhibited an irregular circumference, flat or flaky texture, and a dry, matte appearance. All phenotyping assessments were performed on at least two independent biological replicates to ensure reproducibility. Composite morphotype was determined to be the overall behavior between solid media and liquid media across all tube materials. The composite morphotype is slightly different from the solid media morphotype under certain conditions to account for mixed or non-classical behavior patterns. Specifically, composite morphotype “rough” included strains forming colonies with mixed characteristics on solid media, but noted to have more aggressive aggregation and biofilm formation in liquid media. When solid media morphotype was mixed and the dominant liquid media growth habits ranged from pure planktonic to minimal aggregate or biofilm formation, the composite morphotype for the strain was termed as “mixed.”

### MIC determinations

The minimum inhibitory concentration (MIC) of each drug against individual *Mab* clinical isolates was determined using the standard broth microdilution method (Cynamon et al., 1998; Tilton et al., 1973), based on CLSI guidelines specific to *Mab* (CLSI, 2023). Briefly, powdered drug stocks were obtained from commercial suppliers, except for MRX-6038 (provided by MicuRx Pharmaceuticals) and omadacycline (provided by Paratek Pharmaceuticals). Stock solutions were reconstituted in dimethyl sulfoxide (DMSO), and two-fold serial dilutions were prepared in Middlebrook 7H9 broth to yield final drug concentrations ranging from 128 µg/mL to 0.125 µg/mL for amikacin, amoxicillin, azithromycin, cefdinir, cefditoren, cefoxitin, ceftazidime, cefuroxime, clarithromycin, dicloxacillin, doripenem, faropenem, imipenem, linezolid, metronidazole, moxifloxacin, oxacillin, penicillin V, rifabutin, and tebipenem, and from 16 µg/mL to 0.016 µg/mL for bedaquiline, clofazimine, MRX-6038, and omadacycline. Each well of a 96-well microtiter plate (final volume 200 µL) was inoculated with approximately 10⁵ CFU of *Mab* from an exponentially growing culture. Drug-free *Mab* cultures and broth-only wells were included on each plate as positive and negative controls, respectively. Plates were incubated at 30°C for 72 hours according to CLSI guidelines. Following incubation, bacterial growth was assessed using a Sensititre Manual Viewbox. The MIC was defined as the lowest drug concentration that completely inhibited visible *Mab* growth. All MIC determinations were performed with technical duplicates for each biological replicate. Assays were performed in a minimum of biologic duplicate except for faropenem, tebipenem, metronidazole, penicillin V, cefditoren, MRX-6038, and omadacycline, where supplies were limiting. Additional biologic replicates were performed if there was a greater than two-well discrepancy until a median MIC could be established and reproduced. Median MIC values calculated from the duplicates are reported.

### Mouse infections and *Mab* burden determination

Female C3HeBFe/J (Kramnik) mice (6–8 weeks old; Charles River Laboratories) were used for all experiments as described (Maggioncalda et al., 2020). Each infection was performed at a different time with separate mouse cohorts over a four-year period. Upon arrival, animals were housed under biosafety level 2 (BSL-2) conditions and acclimated for 10–14 days before experimentation. To induce immunosuppression, dexamethasone (Sigma-Aldrich, catalog no. D1756) was administered once daily, seven days per week. Dexamethasone powder (Sigma-Aldrich, Cat. D1756) was weighed for daily dosing and stored at –20 °C in 25 mL polypropylene tubes. Immediately before administration, the powder was reconstituted in 1× PBS (pH 7.4) to the desired final concentration and vortexed at high speed for 2 min, yielding a cloudy white homogeneous suspension. 200 μL of this suspension was administered subcutaneously into the hind dorsal flank using a 27-gauge needle (Becton Dickinson, Cat. 305620). On day 7 after treatment initiation, mice were infected with *Mab* via aerosol using an Inhalation Exposure System (Model A4212, Glas-Col, Terre Haute, IN) equipped with a Collison-type nebulizer. A total of 11 mL of Mab culture grown in Middlebrook 7H9 broth (OD₆₀₀ = 1.0) was nebulized following a standard cycle: 15 min preheating, 30 min nebulization, 30 min cloud decay, and 15 min UV surface decontamination. After exposure, animals were returned to individual cages and maintained under BSL-2 housing.

To determine lung implantation, five mice were euthanized one day post-infection (week 0), and their lungs were collected aseptically. Additional groups (five mice at week 2 and eight per subsequent time point) were sacrificed for organ harvest. Lungs, livers, and spleens were placed in sterile tubes containing 2 mm glass beads and phosphate-buffered saline (PBS, pH 7.4), then homogenized using a Minilys bead beater (Bertin Instruments) for 30 s at 4000 rpm. Homogenates were 10-fold serially diluted in PBS, and 100 μL aliquots were plated on Middlebrook 7H11 agar supplemented with 50 μg/mL carbenicillin (Research Products International, Cat. C46000) and 50 μg/mL cycloheximide (Sigma-Aldrich, Cat. C7698). Plates were incubated for 7 days at 37 °C, and colonies were enumerated. Colony-forming unit (CFU) counts were adjusted for dilution, averaged per sample, and expressed as log₁₀(CFU) for plotting and statistical analysis.

### Histopathology

Lungs, livers, and spleens from mice infected with distinct Mycobacterium abscessus isolates were fixed in 10% neutral buffered formalin for at least 24 hours, then either stored in PBS at 4 °C or submitted directly for processing. Tissues were paraffin-embedded, sectioned at 4–5 μm, and serial slices were stained with hematoxylin and eosin (H&E), Masson trichrome (MAS), and Ziehl-Neelsen (AFB) stains at the Johns Hopkins Hospital Oncology Tissue Services Core. Slides were digitized with unique identifiers using a high-resolution scanner and analyzed in a blinded manner. Histopathologic evaluation focused on the extent and distribution of acid-fast bacilli in tissue, inflammation, collagen deposition, granuloma formation, necrosis, and tissue damage. Representative images and corresponding descriptions were subsequently reassociated with their respective infection groups for data interpretation.

## Data analysis

Raw colony-forming unit (CFU) counts obtained from mouse lung homogenates were compiled and analyzed to determine the mean ± standard deviation for each *Mab* isolate at each sampling time point. Individual data points for all mice were plotted as dot plots to visualize the distribution and variability of *Mab* burdens at each time point.

## Competing Interests

All authors declare no competing or financial interests.

## Author contributions

RAH: experimental design & investigation, methodology, data analysis & interpretation, histopathologic analysis, figure preparation, writing. CT: experimental design & investigation. CMP. methodology and investigation. GC: bioinformatic analysis. BR: methodology and investigation. EC: methodology and investigation NN: methodology and investigation. GL: study conception, study design, investigation, project administration, data interpretation, manuscript preparation, and funding acquisition.

## Funding

This study was supported by NIH grant AI155664. Ruth Howe was supported by the Sherrilyn and Ken Fisher Center for Environmental Infectious Diseases, Division of Infectious Diseases, Johns Hopkins University and by the NIH grant 5T32AI007291-33.

## Acknowledgments

We thank Dr. Charles Daley, National Jewish Health, Colorado, for providing *Mab* clinical isolates M9624, M9626, M9643, M9647, M9654, M9656, M9660, M9672 and M9674, the year they were isolated and their subspecies information. We also thank Dr. Sujan Gautam, Johns Hopkins University, for undertaking some animal studies.

## SUPPLEMENTAL TABLE & FIGURES

**Table S1.** Median MIC values for antibiotics for ATCC 19977 and clinical *Mab* isolates. Abbreviations: CLR clarithromycin; AZM azithromycin; IMI imipenem; DOR doripenem; FRP faropenem; TBP tebipenem; AMK amikacin; CFZ clofazimine; MET metronidazole; LZD linezolid; MOX moxifloxacin; RFB rifabutin; OMC omadacycline; BDQ bedaquiline; PVK penicillin V; AMX amoxacillin; OXA oxacillin; CXM cefuroxime; FOX cefoxitin; CPD cefpodoxime; CDR cefdinir; CAZ ceftazidime.

| Drug Class | Macrolide | Macrolide | Carbapenem | Carbapenem | Carbapenem | Carbapenem | Aminoglycoside | Phenazine | Nitroimidazole | Lincosamide | Fluoroquinolone | rifamycin | tetracycline | Benzoxaborole | Diarylquinoline | Penicillin, 1<sup>st</sup> | Aminopenicillin | Penicillin, 2<sup>nd</sup> | Penicillin, 2<sup>nd</sup> | Cephalosporin, 2<sup>nd</sup> | Cephameycin | Cephalosporin, 3<sup>rd</sup> | Cephalosporin, 3<sup>rd</sup> | Cephalosporin, 3<sup>rd</sup> |
| --- | --- | --- | --- | --- | --- | --- | --- | --- | --- | --- | --- | --- | --- | --- | --- | --- | --- | --- | --- | --- | --- | --- | --- | --- |
|  | CLR | AZM | IMI | DOR | FRP | TBP | AMK | CFZ | MET | LZD | MOX | RFB | OMC | MRX-6038 | BDQ | PVK | AMX | OXA | DCX | CXM | FOX | CPD | CDR | CAZ |
| Isolate |  |  |  |  |  |  |  |  |  |  |  |  |  |  |  |  |  |  |  |  |  |  |  |  |
| ATCC 19977 | 24 | 16 | 20 | 24 | 128 | 128 | 32 | 4 | >128 | 128 | 16 | 32 | 0.5 | 0.25 | 0.095 | >128 | >128 | >128 | >128 | >128 | 64 | 128 | 128 | >128 |
| M9502 | 3 | 24 | 16 | 40 | 128 | 128 | 48 | 4 | >128 | 96 | 20 | 24 | 0.5 | 0.13 | 0.095 | >128 | >128 | >128 | >128 | >128 | 32 | >128 | 48 | >128 |
| M9503 | 8 | 32 | 32 | 40 | 128 | 128 | 24 | 8 | >128 | 96 | 24 | 24 | 0.5 | 0.25 | 0.155 | >128 | >128 | >128 | >128 | 128 | 24 | 128 | 64 | >128 |
| M9507 | 80 | >128 | 32 | 96 | >128 | 128 | 8 | 4 | >128 | >128 | 40 | 12 | 0.5 | 0.13 | 0.07 | >128 | >128 | >128 | >128 | >128 | 48 | >128 | >128 | >128 |
| M9508 | 24 | 128 | 12 | 32 | 128 | 128 | 16 | 4 | >128 | 16 | 16 | 36 | 1 | 0.25 | 0.28 | >128 | >128 | >128 | >128 | >128 | 24 | 128 | 96 | >128 |
| M9510 | 4 | 96 | 32 | 96 | >128 | 128 | 16 | 2 | >128 | 96 | 80 | 12 | 1 | 0.5 | 0.033 | >128 | >128 | >128 | >128 | >128 | 80 | 128 | 96 | >128 |
| M9533 | 40 | 128 | 8 | 24 | 128 | 128 | 48 | 8 | >128 | 96 | 12 | 24 | 0.5 | 0.25 | 0.095 | >128 | >128 | >128 | >128 | >128 | 48 | 128 | 96 | >128 |
| M9624 | >128 | >128 | >128 | >128 | >128 | 128 | 16 | 4 | >128 | 80 | 96 | 3 | 4 | 0.25 | 0.07 | >128 | >128 | >128 | >128 | >128 | >128 | 64 | >128 | >128 |
| M9626 | 24 | >128 | 8 | 48 | 128 | 64 | 24 | 8 | >128 | 80 | 20 | 40 | 1 | 0.25 | 0.095 | >128 | >128 | >128 | >128 | >128 | 80 | >128 | 96 | >128 |
| M9643 | 40 | >128 | 64 | 32 | >128 | >128 | 32 | 4 | >128 | 96 | 16 | 32 | 2 | 0.25 | 0.155 | >128 | >128 | >128 | >128 | >128 | 80 | >128 | 128 | >128 |
| M9647 | 10 | 32 | 64 | 40 | >128 | >128 | 48 | 2 | >128 | >128 | 32 | 32 | 2 | 0.25 | 0.155 | >128 | >128 | >128 | >128 | >128 | 24 | 128 | 80 | >128 |
| M9653 | >128 | >128 | 40 | 96 | 128 | 128 | 80 | 1 | >128 | 12 | 24 | 10 | 0.5 | 0.13 | 0.188 | >128 | >128 | >128 | >128 | >128 | 80 | >128 | 96 | >128 |
| M9654 | 20 | 64 | 32 | 24 | 128 | 64 | 24 | 4 | >128 | 80 | 24 | 24 | 1 | 0.25 | 0.375 | >128 | >128 | >128 | >128 | >128 | 32.5 | 128 | 96 | >128 |
| M9656 | 40 | 64 | 32 | 24 | 128 | 128 | 32 | 4 | >128 | 96 | 40 | 24 | 1 | 0.25 | 0.375 | >128 | >128 | >128 | >128 | >128 | 40 | 128 | 96 | >128 |
| M9660 | 40 | 64 | 4 | 24 | 128 | 64 | 40 | 4 | >128 | 96 | 20 | 12 | 0.5 | 0.25 | 0.19 | >128 | >128 | >128 | >128 | >128 | 48 | 128 | 96 | >128 |
| M9672 | 80 | >128 | 8 | 48 | 128 | 128 | 80 | 2 | >128 | 64 | 16 | 4 | 0.5 | 0.5 | 0.093 | >128 | >128 | >128 | >128 | >128 | 96 | 128 | >128 | >128 |
| M9674 | 2.5 | 8 | 24 | 32 | 128 | 128 | 40 | 2 | >128 | 24 | 8 | 2 | 0.25 | 0.5 | 0.1875 | >128 | >128 | >128 | >128 | >128 | 32 | >128 | 64 | >128 |

**Figure S1.**
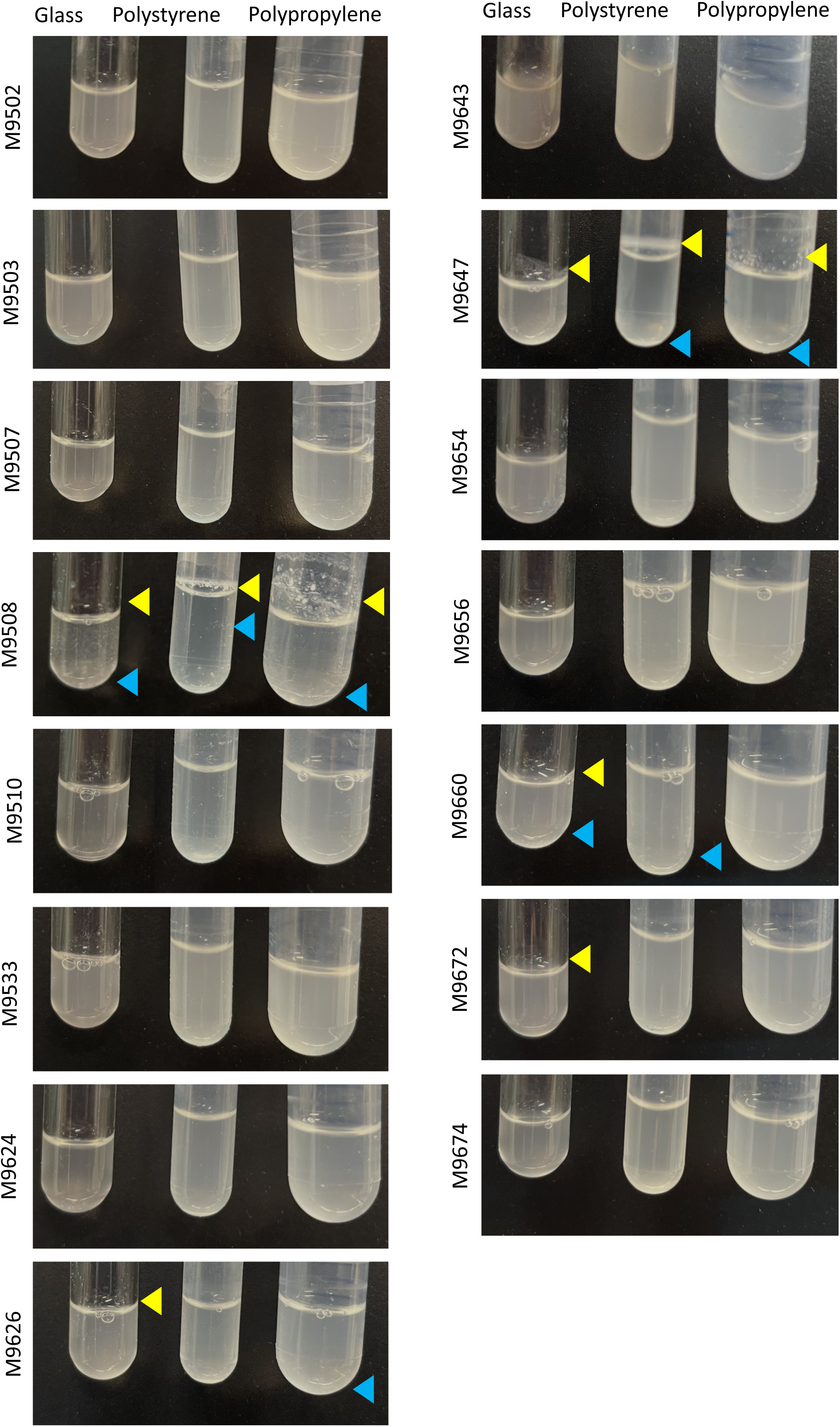
Liquid growth characteristics of pulmonary-derived Mab isolates grown in Middlebrook 7H9 media supplemented with glycerol, ADS, and Tween20 in tubes of different porosity. Yellow arrowheads indicate meniscal biofilm or liminal colony formation. Blue arrowheads indicate non-planktonic aggregates within the liquid media.

**Figure S2.**
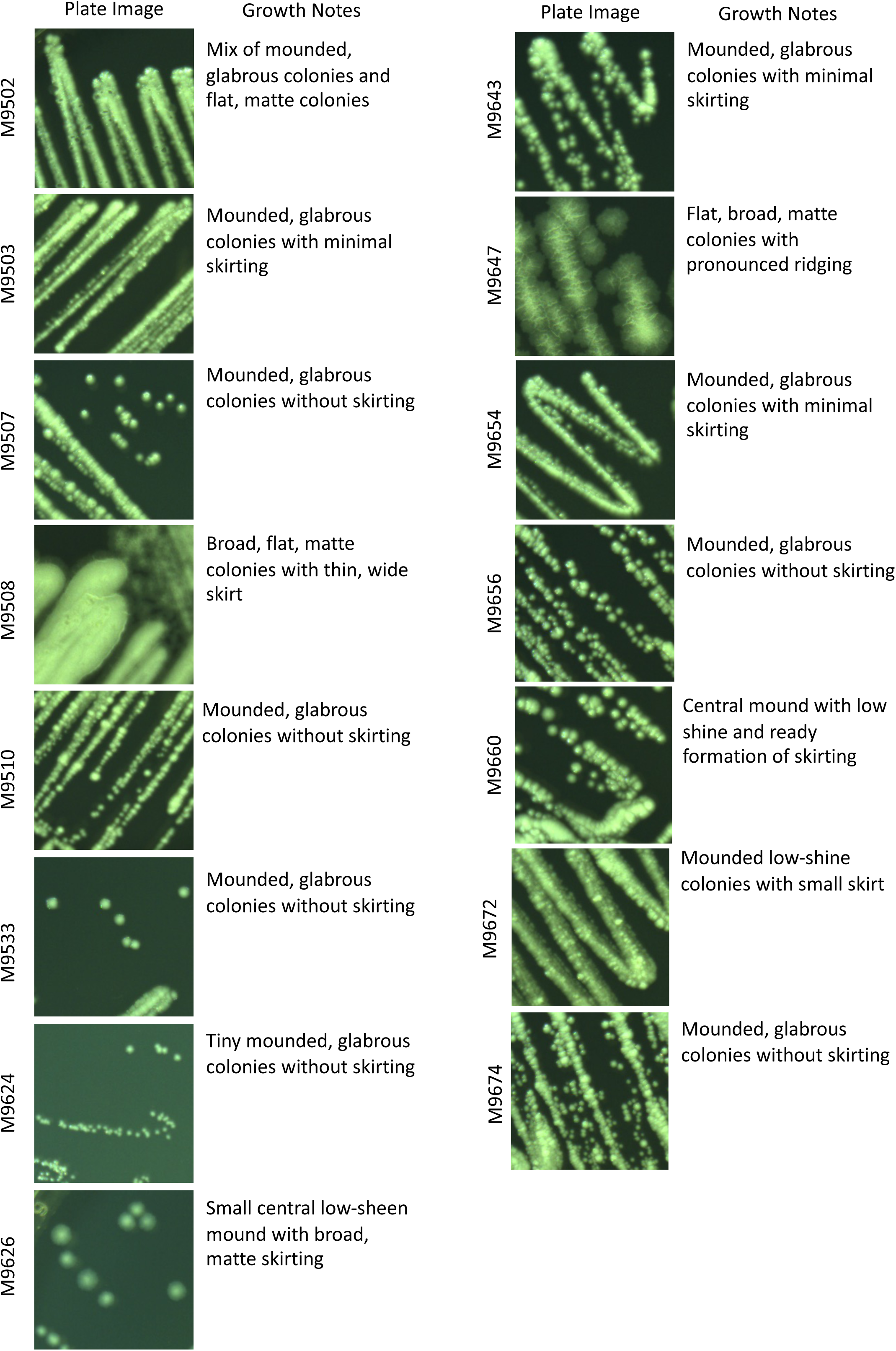
Colony morphologies of pulmonary-derived Mab isolates grown on Middlebrook 7H10 plates supplemented with glycerol and ADS.

## Notes

### Competing Interest Statement

The authors have declared no competing interest.

